# Systematic investigation of interindividual variation of DNA methylation in human whole blood

**DOI:** 10.1101/2024.01.29.577703

**Authors:** Olivia A. Grant, Meena Kumari, Leonard Schalkwyk, Nicolae Radu Zabet

## Abstract

Interindividual genetic variability is well characterised, but we still lack a complete catalogue of loci displaying variable and stable epigenetic patterns. Here, we report a catalogue of stable and variable interindividual DNA methylation in human whole blood by analysing the DNA methylation patterns in 3642 individuals using the IlluminaEPIC array. Our results showed that 41,216 CpGs display stable methylation (SMPs) and 34,972 CpGs display variable methylation levels (VMPs). This catalogue will be a useful resource for interpretation of results when associating epigenetic signals to phenotypes. We observed that SMPs are highly enriched in CpG islands, depleted at CpG shelves and open sea regions of the genome. In addition, we found that the VMPs were under higher genetic control than the SMPs and that trans mQTL pairs are often located in the same TAD or connected by chromatin loops. A subset of these VMPs (784) were classified as putative epialleles and our results demonstrate that these loci located in regulatory regions exhibit a link with gene expression.

## INTRODUCTION

The combination of genetic, epigenetic and environmental variation between individuals is responsible for the large diversity observed in human phenotypes. Large efforts have previously been made to characterise genetic variation in humans, with important advances such as the characterisation of millions of single nucleotide polymoprhisms (SNPs) (1; 2), yet detailed catalogues of epigenetic variation are still not complete.

The need for these efforts are becoming increasingly clear, as research suggests that changes to DNA sequence and exposure to known environmental factors are unable to account for some phenotypic differences that can be observed amongst a population. This has been highlighted by studies in genetically identical organisms, commonly involving mono-zygotic twins but also some inbred animal studies (3). Monozygotic twins in almost all cases, are identical in appearance, however are often discordant for particular diseases or phenotypes. Often, this discordance has been attributed to differences in environmental exposures which can have widespread and significant effects on human health. It is becoming increasingly accepted that epigenetic mechanisms may explain these findings in twin and animal studies. Thus, recent studies have identified epigenetic differences between monozygotic twins who are discordant for particular diseases such as amyotrophic lateral sclerosis, psoriasis and neurofibromatosis, thus suggesting that epigenetic variation could in fact explain differences in phenotype (4; 5; 6).

In understanding the intricate mechanisms underlying epigenetic variation, three main epigenetic marks play pivotal roles: histone modifications, RNAs (such as microRNAs and long non-coding RNAs), and DNA methylation (7). Histone modifications involve the addition of chemical groups to histone proteins which influences the accessibility of genetic material for transcription (8). RNAs, including microRNAs and long non-coding RNAs, are involved in post-transcriptional regulation, impacting gene expression and cellular processes (9). DNA methylation, on the other hand, is a key epigenetic modification involving the addition of methyl groups to cytosine residues in the DNA molecule, often occurring at CpG dinucleotides (10).

Recognizing the significance of understanding the interplay between genetics and epigenetics in shaping phenotypic diversity, our study will focus specifically on DNA methylation. This choice is motivated by the growing body of evidence suggesting that alterations in DNA methylation patterns contribute significantly to phenotypic differences, particularly in instances where genetic and environmental factors alone cannot account for observed disparities (11). By honing in on DNA methylation, our investigation aims to contribute valuable insights into the role of DNA methylation addressing the gaps in our understanding of epigenetic variation.

To this effect, previous studies have aimed to characterise a catalogue of loci showing highly variable DNA methylation in a range of tissues including peripheral blood, cord blood, saliva, placenta and colon (12; 13; 14; 15; 16; 17). Nevertheless, this is a difficult task due to the dynamic nature of the epigenome. Whilst genetic variation is minimal within individuals (intraindividual) and extensive between individuals (interindidvidual), DNA methylation variation is extensive both within and between individuals. This is because individuals within a population can vary for a wide variety of reasons (known examples include stress levels, age, sex and smoking status) all of which in turn, can cause variation in the epigenome. Additionally, these varying patterns may differ between different tissues and cell types, unlike genetic variation.

The Human Epigenome project is an important resource for mapping the human epigenome and some of their efforts were in fact directed towards characterising interindividual variation (18). However these efforts were focused mainly on an approximately 4 Mb region of the genome called the Major Histocompatibility Complex (MHC) rather than genome wide. One important result they found is that DNA methylation variability is often tissue dependent, as the loci they investigated did not show concordant variation across tissues. Furthermore, these highly variable amplicons were mostly intragenic (18), a finding which was also observed in another independent study identifying interindividual Differentially Methylated Regions (DMRs) in monocytes (19). Additional work in human germ cells also supports this idea and further detected a large degree of variation within promoter CpG islands and pericentromeric satellites (20).

A recent study identified variably methylated regions (VMRs), defined as clusters of CpGs with high interindividual epigenetic variation (14). It also found that these regions were enriched at enhancers and 3’UTRs and imprinted loci, which suggests that they may play a functional role in gene expression or biological function.

Additionally, it is now also well known that DNA methylation is strongly influenced by genetic variation. A clear example is provided by the evidence that a rare change at DNMT3L, leads to significant DNA hypomethylation in subtelomeric regions of the genome (21). Nevertheless, DNA methylation can also impact genetic variation and one of the simplest cases is deamination of a methylated cytosine, by which changes a cytosine base gets converted to a thymine base. In this case, the lack of cytosine would result in an absence of DNA methylation, so the variation would result in either no DNA methylation or a fully methylated position (22).

Most evidence showing a relationship between genetic variation and epigenetic variation arises from work characterising the methylation quantitative trait loci, which are genetic variants that influence CpG methylation (23). mQTLs have previously been characterised in brain, blood, lung and adipose tissue (24; 25; 26; 27; 28; 29). Additionally, these methylation quantitative trait loci (mQTLs) may also overlap variants that associate with gene expression levels. Thus, the research linking these epigenetic signatures to genotypes is vital to provide more mechanistic insights into the interplay between genetics, disease and epigenetic variation.

In this study, we identified a robust catalogue of loci within the human genome with either high interindividual variability or high interindividual stability in DNA methylation in whole blood. We do this by leveraging two large datasets from Understanding Society (30) investigating 850,000 CpG sites among a total of 3700 individuals (discovery dataset, n =1175, and validation dataset, n=2570). To our knowledge, this is the largest study using the Illumina EPIC BeadChip (allowing for interrogation of 850,000 sites across the genome) to investigate variability and stability in DNA methylation at CpG sites in whole blood. Other studies focused on the 450K array (17; 14), which has approximately half of the CpGs investigated here or performed bisulfite sequencing which suffers from lower sample size due to higher costs (31; 32).

## RESULTS

### Variably methylated probes and stably methylated probes are widespread across the genome and enriched in regulatory regions

We characterised interindividual variation of DNA methylation in whole blood using a discovery and validation approach using two human sample sets (n=1171 and n=2471, respectively). This analysis was performed on the Illumina EPIC array, which covers 850,000 sites across the human genome. Sites which were known SNP probes, cross hybridizing probes or sex chromosome linked probes were removed from our analysis. Thus, after quality checks and data processing, a total of 747,302 CpGs were included in our analysis (see Material and Methods). We identified 41,216 CpGs to be loci which show robust stable methylation in both our discovery and validation data sets, which we termed stably methylated probes (SMPs), and 34,972 CpGs that display highly variable methylation levels in both our discovery and validation data sets across our population, which we termed variably methylated probes (VMPs) (see Figure 1A and Figure S1A-B; Supplementary File S1 contains the complete list of SMPs and S2 the complete list of VMPs). Approximately 1% of SMPs (553 probes – 3.9 times more than expected by chance, Fisher’s exact test p-value = 5 *×* 10^−4^) are in non-CpG context which are expected to be unmethylated and show no interindividual variation in methylation, as we would expect in non stem cells (33). Half of the VMPs and 20% of the SMPs are EPIC specific, which means they could not have been previously identified on studies that considered only 450K array datasets (Figure S1C-D). Interestingly, SMPs showed either low or high methylation, whereas VMPs showed predominantly intermediate methylation levels (see Figure S2A-B). Given that interindividual variation in DNA methylation may relate to environmental differences, we investigated the overlap of VMPs and SMPs with CpGs associated with phenotypes that are known to have strong associated epigenetic signatures, namely: age, smoking and sex. We found that despite VMPs having a higher overlap with known sex, smoking or age associated CpGs, than SMPs, the variation in methylation for the majority of VMPs cannot be explained by these phenotypes alone, indicating that they are under the control of other environmental or genetic factors(see Figure S2C-D).

**Figure 1.**
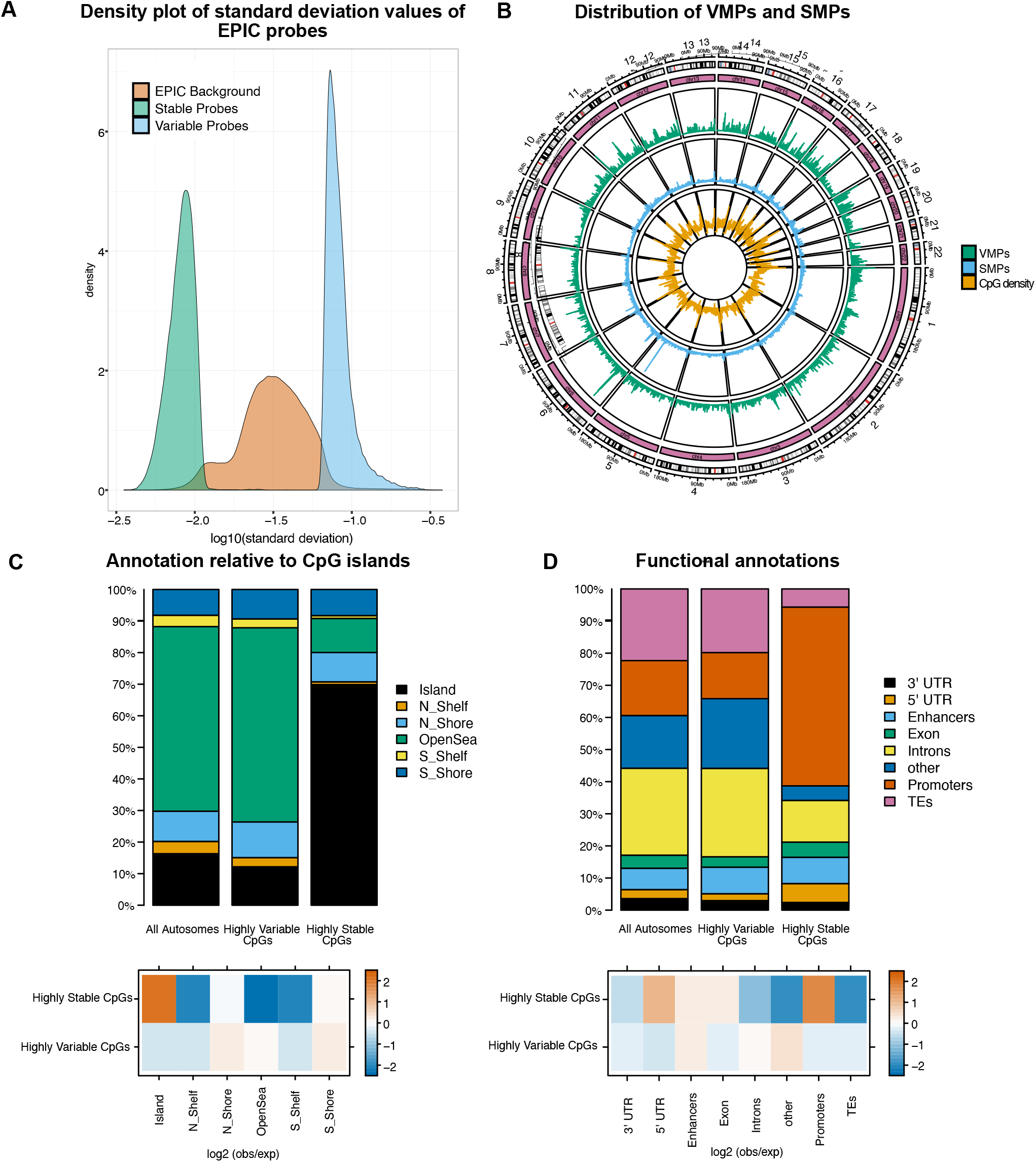
Identification and annotation of variably methylated probes (VMPs) and stably methylated probes (SMPs). (A) Density plot representing the standard deviation values of methylation across our samples for VMPs, SMPs and the autosomal EPIC background. (B) Circos plot representing distribution of: CpG island density, SMPs and VMPs across the genome (at 50,000 Kb resolution). (C) The annotation of VMPs (n= 34972) and SMPs (n=40288) relative to CpG islands compared to the autosomal background. Bottom panel shows the log2 (obs/exp) annotations based on the autosomal EPIC background of the different annotations. (D) The annotation of VMPs (n= 34972) and SMPs (n=40288) to genomic features compared to the autosomal background. Bottom panel shows the log2 (obs/exp) annotations based on the autosomal EPIC background of the different annotations. Note that all autosomes include the EPIC background, VMPs and SMPs, with the majority (90%) consisting of the EPIC background.

Blood cell type composition varies among individuals, and each cell type has a distinctive DNA methylation profile, so this is one potential cause of DNA methylation variation. The effect for an individual CpG is an average across cell types of the product of the abundance difference and the DNA methylation difference. Indications are that this is not large enough to drive the VMPs that we observe. Although we did not correct or deconvolute the data for cell composition, only 42 of the 450 CpGs used to distinguish cell types by *flowsorted*.*blood*.*epic* (34) overlap with our VMPs and none with our SMPs. Those 450 probes do not consist of all of the cell type differences, but they were chosen as optimal for distinguish cell types and so one would expect that majority of those probes are in the VMP list if cell type is an important driver of VMPs.

To identify possible biological pathways that these probes were involved in, we performed KEGG ontology analyses which showed several enriched terms for these VMPs and SMPs, supplementary Table S1 shows the top enriched terms for VMPs and SMPs. VMPs were enriched for terms such as Neuroactive ligand receptor interaction, olfactory transduction, cushing syndrome and morphine addiction. Enriched terms for the SMPs revealed several signalling pathways, such as metabolic pathways, MAPK, HIF-1 and FoxO signalling pathways along with other terms such as oxidative phosphorylation and lysine degradation.

Next, we investigated if the VMPs and SMPs are assigned to either housekeeping or tissue specific genes. Unsurprisingly, our results show that SMPs are usually enriched at housekeeping genes, while VMPs at tissue specific genes (see Figure S2E).

Both VMPs and SMPs are distributed evenly across the genome, but SMPs are concentrated in CG dense regions (Spearman correlation coefficient of 0.36 between SMPs and CpG density and 0.07 between VMPs and CpG density); see Figure 1B. To help us gain more insight into the functional role of these variably and stably methylated probes, we characterised their location with respect to CpG islands and functional regions. We observed that SMPs were highly enriched in CpG islands and depleted at CpG shelves and open sea regions of the genome. In contrast, we found that VMPs were depleted at CpG islands and enriched at CpG shores, but also depleted at CpG shelves (see Figure 1C). Furthermore, SMPs were also highly enriched at 5’ UTRs and promoter regions of the genome, and showed slight enrichment at enhancers suggesting that they are unmethylated. On the other hand, VMPs were enriched at intergenic regions, introns and enhancers, but depleted at UTRs, promoters and transposable elements (see Figure 1D). Altogether we found that SMPs are located at housekeeping promoters and VMPs are located in regulatory regions (enhancers and intergenic regions) of tissue specific genes.

Imprinted control regions display high methylation on either maternal or paternal copies of the DNA with the other copy being unmethylated (35). Given the specific signatures of methylation at these imprinted regions, we also investigated whether variable probes were enriched at imprinted regions of the genome. While only one single stably methylated probe overlaps imprinted regions, we found 242 VMPs that overlap an imprinted gene, which is higher than we would expect by chance compared to the EPIC array background (Permutation test, p-value 0.001); see Figure S2F.

### VMPs and SMPs located at promoters and enhancers are enriched for transcription factor motifs

DNA methylation has been shown to impact binding of TFs to DNA (36; 37), which downsteam can impact gene regulation. To investigate this, we next performed transcription factor (TF) binding motif enrichment and gene ontology analyses for our VMPs and SMPs located at promoters and enhancers specifically. First, we evaluated whether the VMPs and SMPs located at promoters were enriched in motifs for TFs (50 bp window). For the VMPs we found 408 unique enriched TFs meaning these 408 were specific to VMPs only, with strongest evidence for TFAP2A and NHLH1 (Figure 2A). For SMPs, we identified 86 unique enriched TFs (specific only to SMPs), including SREBF1 and AHR (Figure 2A). Similarly, for VMPs and SMPs located at enhancers, we identified 376 uniquely enriched TF motifs at VMPs and 74 enriched at SMPs (Figure S3A). The most strongly enriched TFs at enhancers differed to those at promoters, with ATF2 and FOS at VMPs and NNT and ODC1 at SMPs (see Figure S3A).

**Figure 2.**
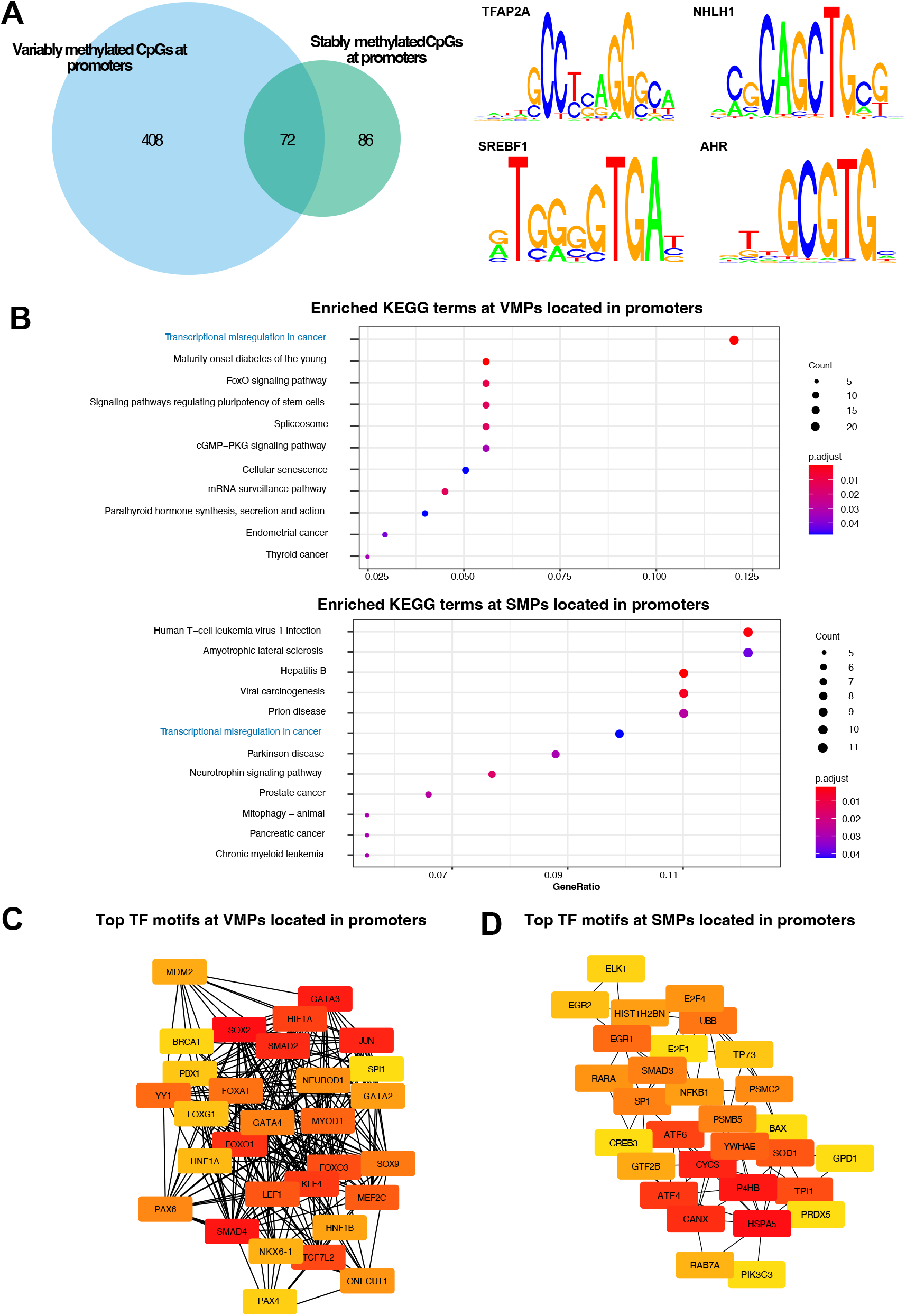
Transcription factor motif enrichment for VMPs and SMPs at promoters. (A) Overlap of enriched TF motifs for VMPs (blue) and SMPs (green). The top two motifs enriched at VMPs were TFAP2A and NHLH1 and at SMPs were SREBF1 and AHR. (B) KEGG analyses for the significantly enriched TF motifs at VMPs and SMPs. Common terms are highlighted with blue colour. (C-D) Sub networks of the top 30 enriched TF motifs at (C) VMPs and (D) SMPs. Node colour represents the degree of connectivity. The scale from red to yellow represents the top 30 enriched TF motif rank from 1-30, with red indicating highest degree and yellow indicating lowest degree.

To investigate whether the transcription factor motifs were enriched for terms related to biological processes or pathways, we performed pathway analyses using the GO and KEGG databases. We identified several enriched KEGG pathways for the TFBS enriched at VMPs in promoters, spanning a wide range of processes such as transcriptional misregulation in cancer, FoxO signalling pathways and signalling pathways regulating pluripotency of stem cells (see Figure 2B). We also identified several enriched terms for the TF motifs enriched at SMPs in promoters such as viral carcinogenesis, prion disease and Parkinson disease (see Figure 2B). Interestingly, for those TF motifs enriched at VMPs in enhancers, we find similar pathways, such as transcriptional misregulation in cancer (see Figure S3B).

We then used cytohubba (a cytoscape tool) to identify if any of these transcription factors are regulatory hubs controlling many genes. For the TF motifs enriched at VMPs within promoters we identified MAPK1, GATA3 and SOX2 and other genes to be hub TFs in the network (see Figure 2C). Moreover, for those TFs enriched at SMPs within promoters we identified CYCS, P4HB and HSPA5 to be hubs in this network (see Figure 2D). Similar to our enrichment analyses, we also found a large overlap between the hub genes identified at promoters and enhancers for both VMPs and SMPs. This suggests that enrichment of transcription factors differs slightly according to functional regions in the genome, but not significantly and that the hub transcription factors enriched near highly variable and highly stable methylation sites are robust. Interestingly, both VMPs within enhancers and promoters display enrichment of motifs for JUN, HIF1A and FOS. TET enzymes which are involved in active demethylation, display sequence preferences for these motifs, which indicates a potential mechanistic link between the enrichment of these TFs at VMPs and changes in DNA methylation (38).

### Approximately half of the VMPs are under genetic control

To gain insight into the mechanism that can explain the variability or stability at these CpGs, we considered genetic cases and, thus, analysed a whole blood mQTL dataset (39). Of the VMPs, 44.9% were associated with SNPs (see Figure 3A). We further categorised these into *cis* and *trans* mQTLs where a *cis* mQTL was defined when the SNP and CpG were less than or equal to 500bp away from one another, a *trans* mQTL is defined when the SNP and CpG were more than 500bp away from one another. The intention in using such a small window (500 bp) was to distinguish differences driven by sequence variation in the CpG, the probe sequence or immediately flanking sequences at nucleosome scale. Of the VMP mQTLs, 21% were *cis*, 79% were *trans* mQTLs. One example is an mQTL pair located on chromosome 1 at the probe cg04315214 which is associated with two independent SNPs at the gene PRKCZ (FDR <0.01). For SMPs, only 3.27% of the SMPs were associated with SNPs (see Figure 3B), with 8.1% being *cis*, and 91.9% *trans*. An example of an SMP mQTL is on chromosome 1 between cg04093404 and 4 SNPs at the gene TAS1R1. These results indicate that a large part of variability in DNA methylation could potentially be explained by genetic variation, while genetic variation does not seem to be linked to stability of methylation status (methylated or unmethylated cytosines).

**Figure 3.**
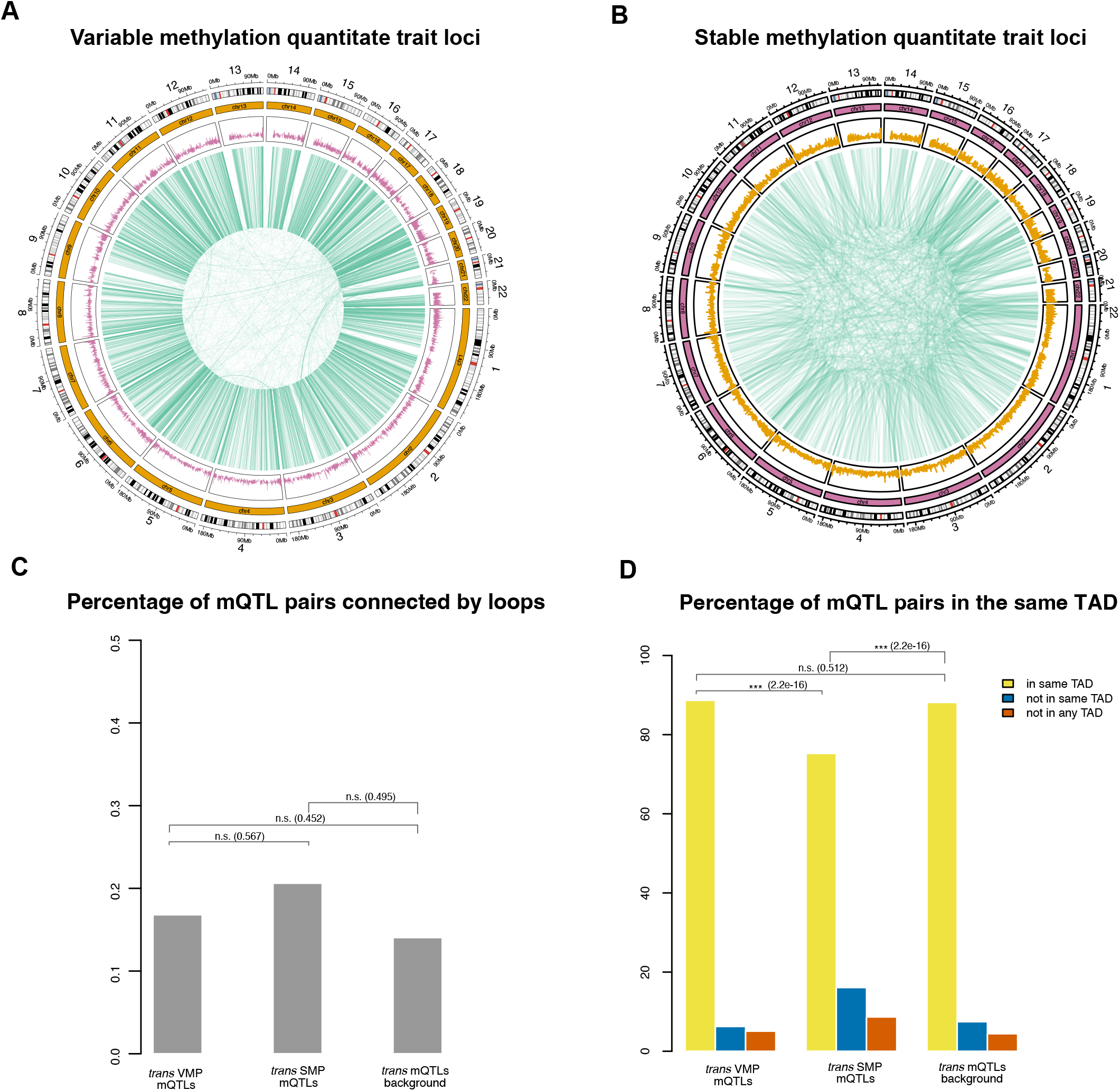
Methylation quantitative trait loci connect VMPs and SMPs with SNPs. (A-B) Circos plot illustrating *cis* and *trans* mQTLs in whole blood. The outermost ring displays chromosome numbers and bands. The second ring represents CpG island density. The third ring represents associations between (A) VMPs and SNPs and (B) SMPs and SNPs (green lines).(C) Barplot representing the percentage of *trans* mQTL pairs connected by chromatin loops. (D) Barplot representing the percentage of *trans* mQTL pairs in the same topological associated domain. Fisher’s exact test was performed to determine statistical significance against the Illumina EPIC array background (p-value: n.s. *≥*0.05, ^*^ < 0.05, ^**^ <0.01 and ^***^ <0.001).

We also tested whether our SNP-CpG pairs were enriched in regulatory regions such as promoters or enhancers, however we found no significant enrichment in any particular regions that we would expect by chance. Mechanistically, the *trans* mQTLs could be explained by the 3D chromatin organisation of the DNA, with SNPs and CpG sites that are distal on 1D, making 3D contacts. Thus, we hypothesised that the *trans* SNP-CpG pairs may occur within topologically associated domains (TADs) and at chromatin loops as genetic and epigenetic interactions.

Figure 3C shows that the trans mQTL-SNP pairs were annotated to a very low percentage of loops for VMPs, SMPs and EPIC background. Chromatin loops call very strong interactions, but TADs identify regions that interact more inside than outside thus allowing to capture finer connections between SNPs and VMPs. Our results show that a higher percentage of VMP mQTL-SNP pairs occupy the same TAD than SMP mQTL-SNP pairs and than we would expect by chance alone (Fisher’s exact test p-value <0.05). Several of the mQTLs overlapped multiple genes, including genes such as DNMT1, SEPTIN9 and ILF3 (Figure 4). Enrichment analyses of VMP-mQTL associated SNPs revealed several GO terms such as cell morphogenesis involved in neuron differentiation, small GTPase mediated signal transduction and cation transmembrane transporter activity (see Figure S4A). These SNPs were also enriched for few KEGG terms including Focal adhesion, axon guidance and cell ahesion molecules (see Figure S4C). The SMP-mQTL associated SNPs were enriched for fewer GO terms, but included pathways such as nuclear chromosome part, response to peptide and chromatin (see Figure S4B). However, were only enriched for one KEGG term, AMPK signalling pathway (see Figure S4D).

**Figure 4.**
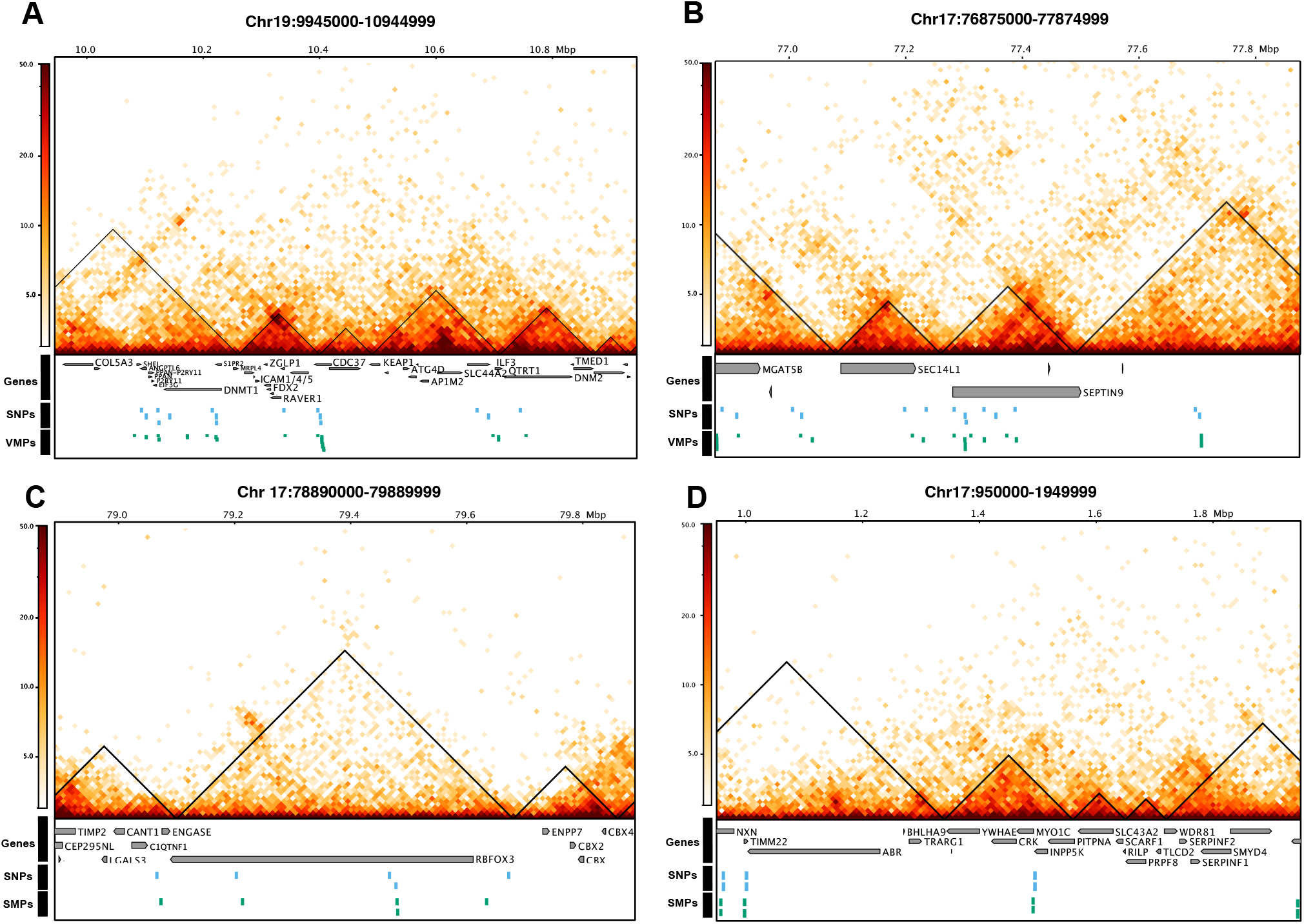
mQTL pairs occupy the same topologically associated domains. (A) and (B) illustrate cases where VMPs and their associated SNPs occupy the same TADs (C) and (D) illustrate cases where SMPs and their associated SNPs occupy the same TADs. x-axis indicates genomic position, y-axis represents the distance between the two genomic regions and the colour the number of contacts between the two genomic regions (with darker colours indicating more contacts retrieved by HiC experiment and lighter colours representing fewer contacts). Black lines indicated TADs, green boxes indicate VMPs/ SMPs, blue boxes indicate SNPs and grey boxes indicate Genes which are appropriately labelled.

### DNA methylation variation in 5’UTRs at putative epialleles is linked to gene expression variation

*Epiallele* is a term that has been defined variously and often in negative terms (40), that is as sites which are variably expressed but not alleles, or even more strictly in the absence of a genetic difference (41). Here, in contrast we screen for *loci* which are present in more than one epigenetic state, which could be stochastic or driven by genetics, environment or parent of origin, all falling under a broad definition of *epiallele*. Some of the variable methylation probes display two peaks, with one subgroup of the population displaying lower methylation at a CpG site and the other subgroup high methylation, which indicates these are potential epialleles (Figure 5A). To investigate this sytematically, we screened for putative epialleles in human whole blood by applying a test for unimodality (using the Hartigan’s dip test) in CpG sites which showed a variable intermediate methylation value. A total of 784 CpG sites met this criterion and we labeled them as putative epialleles (see Figure 5B and Supplementary File S3). 405 of these putative epialleles are associated with at least one gene, and 52 of these genes contained more than one putative epiallele, with PM20D1 containing 8 putative epiallele. PM20D1 is a gene which has previously been reported to be associated with obesity, insulin resistance and the progression of Alzheimer’s disease (42; 43). Moreover, it has previously been reported to contain a variably methylated region associated also with body mass index (44). Additionally, several genes involved in the major histocompatibility complex such as HLA-DRB1, HLA-DQA1, HLA-C and HLA-DRB6 also contained several CpGs identified as putative epialleles in our analysis. Of these CpGs, 42% are found on the Illumina 450K array and 58% were unique to the Illumina EPIC array (see Figure 5C). Furthermore, we hypothesised that some of these putative epiallele sites may be controlled by genetic variation and found that roughly 19% of these were in fact not linked to any genetic variants, but 81% were. Of these, the majority were *trans* mQTLs (65%) and the others were *cis* mQTLs (16%) (see Figure 5D).

**Figure 5.**
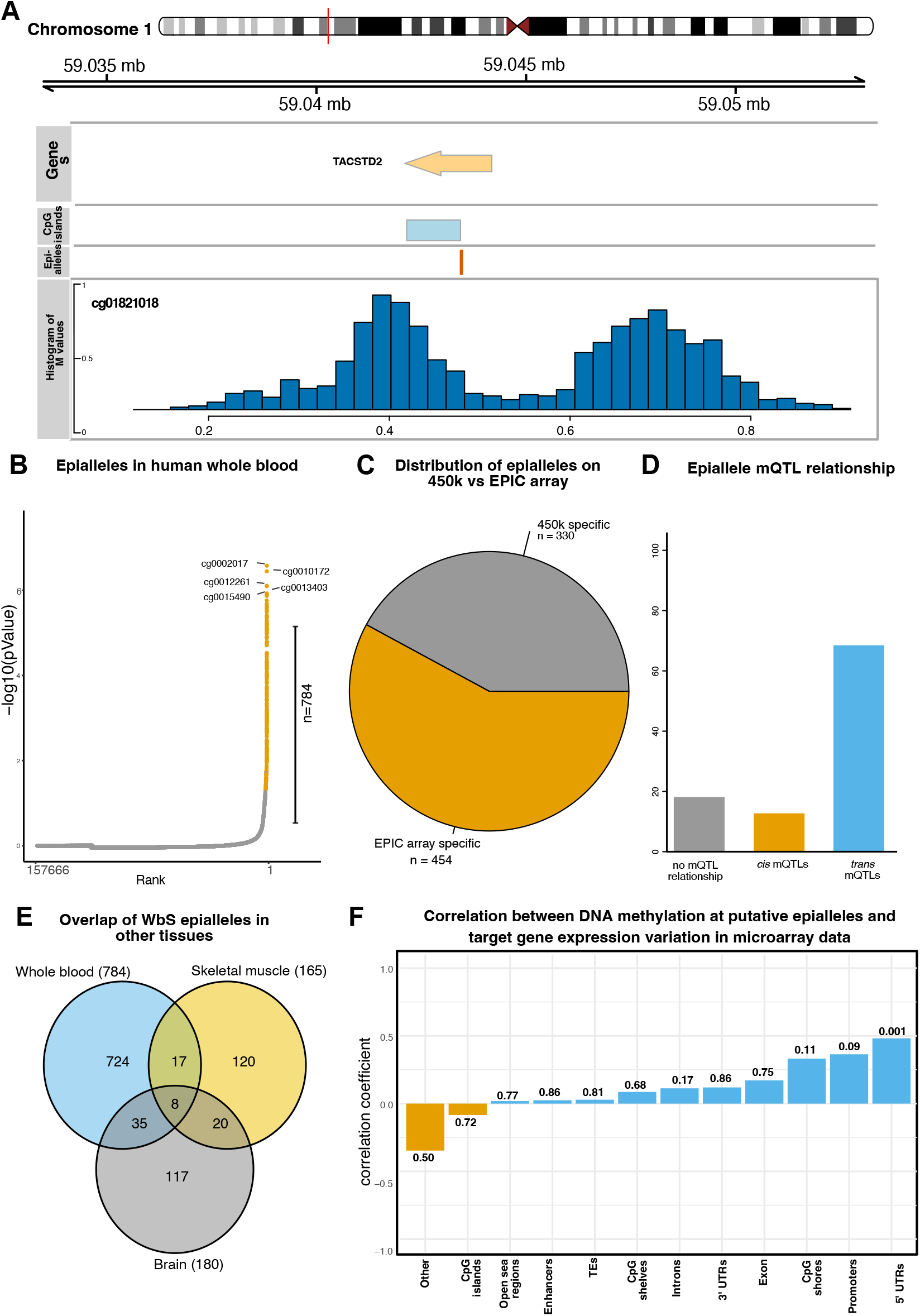
Putative epialleles identified in whole human blood. (A) Genomic plot showing a putative epiallele on chromosome 1 overlapping the TACSTD2 gene. (B) Rank plot showing proportion of CpG sites on the EPIC array which are broad-sense epialleles by Hartigan’s dip test. A total of 784 CpGs met this threshold, and the top 5 are annotated. (C) Pie chart showing distribution of putative epialleles on the Illumina 450k and Illumina EPIC array. Barplot showing percentage of mQTL relationship of putative putative epialleles. Y axis represents percentage. Venn diagram indicating overlap of putative putative epialleles identified in whole blood, brain and skeletal muscle tissue in humans. (F) The correlation between variation in gene expression and variation in methylation of epialleles. The correlation test significance is reported above the bars (p-value: n.s. *≥* 0.05, ^*^p-value<0.05, ^**^<0.01 and ^***^ <0.001).

As we identified that some of these epiallele sites were potentially mediated by genetic variants, we aimed to investigate if these are tissue specific or common between different tissues. Therefore, we calculated these putative epialleles in two other human tissues from different cohorts; first, 160 skeletal muscle samples (45) and secondly, 1121 brain samples (46) using the same method explained in *Materials and Methods*. We identified limited overlap between all three tissues (Figure 5E), suggesting that the majority of these putative epialleles are tissue specific. These putative epialleles are also located within tissue specific genes, which further supports the result that they are tissue specific (Figure S5A). In addition, 7 of them are located at imprinted loci, including OR2L13 (cg08861456), RGS14 (cg16006841), WDR27 (cg11938672 and cg18322025), BRSK2 (cg08208480), ZNF331 (cg05338009) and GNAS (cg24617313) (Figure S5B).

To try to investigate the regulatory role for these epialleles, we also checked their functional annotations. Interestingly, we find the putative epilleles to be enriched at enhancer and intergenic regions, but depleted at promoters, exonic and 3’UTR regions (Figure S5C-D). One possibility is that these epialleles are under control of Polycomb Repressive Complex and, to test that, we considered the co-localisation of epialleles with H3K27me3 (a epigenetic mark deposited by PRC2) for different blood cell lines from ENCODE project (47). Our analysis showed that only small percentage of epialleles (less than 10%) are located at H3K27me3 ChIP-seq peaks (Figure S5E).

As it has previously been reported that epigenetic variation at epialleles may result in gene expression variation, we next annotated putative epialleles to their nearest target gene in order to investigate this further. We calculated the relationship between methylation variation and target gene expression variation in microarray data using the Pearson’s correlation. Our results show a positive correlation between methylation variation at putative epialleles in 5’ UTRs, promoters and CpG shores and their target gene expression variation (Figure 5F). It is worthwhile noting that we were able to find a statistically significant link between methylation variation of epialleles located at 5’UTRs and gene expression variation despite not using matched expression and methylation datasets. This raises the question whether VMPs in general, rather than putative epialleles, can impact gene expression since they are also enriched at regulatory regions and for TF motifs. We performed the same analysis as for putative epialleles, but in this case we found negligible correlation (Pearson correlation coefficient lower than 0.15) between variability in gene expression and variability in DNA methylation at VMPs (Figure S6).

## DISCUSSION

We analysed DNA methylation interindividual variability and stability amongst healthy individuals in a cohort of 3642 individuals from UK (Understanding Society datasets) in order to better understand its role in diversity in human phenotypes and its relationship with gene expression variation and other genomic factors. We were able to establish the presence of epigenetic interindividual variation (VMPs) and stability (SMPs) across the human genome and showed that these loci were widespread across the genome. When comparing the VMPs discovered in this study to findings previously reported in other studies, we do recover some of the previously reported VMPs, but also identify novel ones. For example, in our VMPs, we recover 85.28% of the hyper variable CpGs identified by (17) in whole blood that are also found in other tissues, 43% of previously reported VMPs in different blood cell types from 426 monozygotic twin pairs (14), and 41% of the CpGs from the EPIC array that are located in regions displaying variability in methylation in multiple tissues from (31; 32) (Figure S7).

Overall, we recover more CpGs sites that display interindividual variability in DNA methylation (approximately 31K additional CpGs) and there are two reasons for this. First, our study is the first one to our knowledge that uses the EPIC array to investigate interindividual variation in DNA methylation and approximately half of our VMPs are EPIC array specific (Figure S1C). Second, these previous studies investigated CpGs that display interindividual variation in DNA methylation in multiple tissues. When applying our same approach to EPIC data in brain (46) (Supplementary File S4) and skeletal muscle (45) (Supplementary File S5), we found that that 5835 VMPs are common between whole blood and brain or skeletal muscle. This is similar to the number of CpGs that display interindividual variation in DNA methylation in multiple tissues in previous studies (17).

Our analysis is rooted in a population that is representative, avoiding biases towards specific disease states or aging. Nevertheless, we identified a relatively small subset of our VMPs that is associated with age-related CpGs. We anticipate that running a similar analysis on datasets curated specifically for aging populations (e.g., 48) would likely reveal an increased number of VMPs associated with age. Future research could delve deeper into age-related changes in DNA methylation by focusing on aging populations, identifying additional layers of complexity in epigenetic regulation. While our study presents a comprehensive analysis, we acknowledge the need for targeted investigations to explore age-related nuances and specific health conditions in more depth.

It is important to note that most of population large scale studies are performed in whole blood and, recently, EPIC array become the most used assay for Epigenome Wide Association Studies (70% of EWAS studies deposited on GEO used EPIC array in 2020) as WGBS is prohibitively expensive for large population studies (49). This is exactly what our study focuses on and having this catalogue of highly variable probes would be of great use when investigating epigenetic signals with EPIC array in whole blood.

Overall, our study indicates that approximately 35K CpGs vary in a meaningful way on the EPIC array in whole blood. Assaying only those that vary meaningfully could significantly increase the statistical power of an EWAS. However, we do not advocate focusing only on these, since one might miss probes that display variability in certain conditions, disease states or upon exposure to certain environment. Nevertheless, this catalogue of probes can provide additional information when interpreting results from EWAS studies, especially for whole blood.

Most importantly, in this study, we have performed a systematic characterisation of the VMPs and SMPs to identify potential underlying processes that could explain the interindividual variation in DNA methylation. Consistent with previous research (50), we demonstrated that only a small percentage of these loci overlap regions known to be associated with biological age, biological sex and smoking status. Interestingly, we observed that all of the highly variable sites showed an intermediate methylation status, a finding previously identified by (13). The role of intermediate methylation remains unclear, however, previous work suggests that it may be a conserved signature of gene regulation and exon usage (51). Regions of intermediate methylation were also shown to have similar intermediate levels of active chromatin marks and their target genes also having intermediate transcriptional activity. This together with the enrichment of these sites at CpG shores, intergenic regions and enhancers, which was also previously reported in (14), indicate that these sites may have regulatory potential distinct from repressive or permissive states resulting from fully methylated or unmethylated sites. In contrast, the most stable sites show either high or low average DNA methylation values across our cohort, indicating these loci might be under tight epigenetic control. In line with this, we identified an enrichment of SMPs at housekeeping genes in contrast to the enrichment of VMPs at tissue specific genes.

Expanding on this, it has also been reported that highly variable sites are found at imprinted control regions (17). Through investigation of the genes annotated to our VMPs, we also found 21 VMPs identified in our analysis annotated to the gene HOXA5, a gene predicted to be maternally imprinted. We therefore checked the enrichment of our identified VMPs and SMPs in imprinted regions, and found that the VMPs were significantly enriched in imprinted regions compared to SMPs (permutation test, p-value <0.05). This is also in line with previously reported similar results (17; 14; 52).

The relationship between DNA methylation and gene expression is a complex one. For example, it is traditionally thought that DNA methylation at promoter regions is linked directly to transcriptional repression or gene silencing. Despite this, recent work that synthetically methylated thousands of promoters in the human genome failed to identify a link between promoter DNA methylation and gene expression. Yet, they suggest that the context specific roles of DNA methylation are highly influenced by transcription factor binding affinities (53). Therefore, we investigate the regulatory networks by searching for the presence of transcription factors binding sites among VMPs and SMPs enriched at important regulatory regions such as promoters and enhancers. Interestingly, we identify the presence of transcription factors which have indeed been previously reported to be methylation sensitive. For example, we identify TFAP2A as the most signficantly enriched motif at VMPs at promoters. It has previously been reported that the presence of DNA methylation leads to an increased binding of TFAP2A to B1, leading to suppressed gene expression of NRBP1 gene (54). Additionally, we find SREBF1 to be the most significantly enriched TF motif at SMPs at promoters which has also been reported to be sensitive to CpG methylation (55). Furthermore, our hub analysis for transcription factor motif at VMPs and SMPs located in enhancers and promoters also revealed more methylation sensitive TFs such as JUN, ATF4 BRCA1, FOXA1 and HIF1A (56; 57; 54; 58). This indicates that interindividual differences in DNA methylation would result in differences in TF binding and consequently differences in gene regulation.

It is possible that variance or stability of DNA methylation may also be directed by genetic differences, as previously reported (24). In line with this, we showed here that VMPs seem to be under higher genetic control than SMPs, suggesting that genetic differences may in part drive epigenetic interindividual variability. While there are several ways that genetic differences may result in epigenetic differences in *cis*, the mechanism by which mQTL pairs work in *trans* remains unclear. We therefore hypothesised that 3D chromatin organisation and DNA methylation are tightly linked based on previous work indicating that TADs play a role in gene regulation (59). With this aim, we found that majority of *trans* VMP mQTL pairs were located in the same TAD, but this was not significant as it is not more than we would expect by chance too (see Figure 3D). This indicates that mQTL pairs (both for VMP mQTLs and non VMP mQTLs) are in part influenced by or influence chromatin organisation, suggesting that 3D genome organisation may in part explain the interindividual variability in DNA methylation (see Figure 3D). However, it is important to note that although the design of the Illumina EPIC array was curated to include more distal regulatory elements such as enhancers, it is possible that the design of the array was not ideal to answer this research question.

Epialleles are regions at which the epigenetic state varies amongst individuals within a population (60). DNA methylation at epialleles occurs during embryonic development and leads to prominent interindividual variation (61). Furthermore, intermediate methylation states have previously been reported as a feature of epialleles in humans (62) with a bimodal distribution of either very high or low methylation. Therefore, we leveraged our catalogue of VMPs as a means of identifying putative epialleles as described in Materials and Methods. Using this approach, we were able to identify 784 putative epialleles in human whole blood which showed intermediate methylation levels across our sample, yet upon closer inspection demonstrated bimodal distribution, with either low or high methylation within individuals. We observed that just over half of these are unique to the Illumina EPIC array, meaning we were able to reveal novel putative epialleles. The establishment of these putative epialleles remains unclear, but may result from several factors. One possibility is that the differences in methylation state at these sites is due to genotype. Whilst we screened out CpGs related to SNPs, some epiallele sites may still be influenced by mQTLs. Here, we were able to identify enrichment for mQTLs at putative epialleles and these were mainly *trans* relationships. However, there were still a portion of putative epialleles which were not linked to any SNPs via our mQTL analysis, suggesting that stochastic or environmental exposures may play a role in the establishment of these putative epialleles in whole blood. We further demonstrated that some of these putative epialleles display tissue specificity (Figure 5E).

Epiallelic variation is functionally characterised in terms of its influence over gene transcription (60), however, the direction of effect remains debated in the literature. For example, one study suggests that differential epigenetic modification of two distinct variants is associated with tissue-specific changes in adjacent gene expression (62). Therefore, we explored this idea further by focusing on the relationship between DNA methylation variance and target gene expression variance at these putative epialleles and also by considered the genomic context in which DNA methylation was found, to try and unravel this relationship further. Interestingly, it is observed that DNA methylation variance and target gene expression variance are positively correlated when epialleles are located at 5’ UTRs, promoters and CpG shores. This identifies a link between DNA methylation at specific regulatory regions and gene regulation. It is important to note, that this analysis was performed in non matching data sets (i.e. the data sets were collected from individual cohorts), which further supports the generalisation of these results. More importantly, this link between DNA methylation and gene expression seems to be specific for this putative epialleles, given that a similar analysis over all VMPs was not able to identify similar correlations with gene expression.

The identification of putative epialleles that are linked to gene expression and are susceptible to genetic and environmental exposures may be indicative of an adaptive mechanism as previously described (63; 62), making these sites important for research exploring adaptive responses to environmental exposures.

## MATERIAL AND METHODS

### DATASETS AND STATISTICAL ANALYSIS

In this study, we used whole blood Illumina Infinium MethylationEPIC BeadChip DNA methylation data collected from participants involved in Understanding Society: The UK Household Longitudinal Study; see (64; 65). We chose these data sets so that we were able to perform our analysis on relatively homogenous populations using a discovery and validation approach. Furthermore, this data was not collected for any various specific exposures such as ageing, smoking behaviour or any health condition, meaning this data would be a representative dataset for the British population. Thus, the VMPs that we identified in this study are not biased by any exposure or health condition. Following quality checks of the data, our final data set consisted of 1171 participants for discovery and 2471 participants for validation. Raw signal intensities were processed using the R package bigmelon (66) from IDAT files and used same preprocessing steps as in (65). Briefly, the data was then normalised via the interpolatedXY adjusted dasen method implemented in the R package, watermelon (67). Following normalisation of the data, SNP probes, cross hybridizing probes (68) and X or Y linked probes were removed from the data set. Moreover, as sex of our samples was self reported, we also utilised a DNA methylation based sex classifier (69) in order to remove samples where reported and biological sex did not match. This resulted in 4 and 11 samples being removed from our discovery and validation data sets, respectively. Lastly, to avoid the possibility that this analysis would flag up SNPs from our sample, we looked for confounding between SNPs and CpG sites using the MethylToSNP R package (70). The function *MethylToSNP* extracts CpG sites corresponding to genotypes CT,TT and CC using 3 discrete levels of methylation: fully methylated, fully unmethylated, and 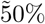 methylation. This resulted in the identification of CpG sites confounding with SNPs and these were also removed from our analysis. The final discovery and validation data set consisted of 1171 and 2471 samples respectively, and 747,302 DNA methylation sites.

Two previous studies use analysis of duplicate arrays to classify probes by reliability on the basis of an interclass correlation (ICC) threshold (71; 72), which we do not think is suitable as a filter. Duplicates in principle allow estimation of biological (between individuals) variance and technical (between duplicates) variance. Probes may differ in their reliability (technical variance), but it cannot be usefully compared to the biological variation because, in a given, sample there may often be little or no true biological variation. Low ICC probes can equally well be interpreted as invariant (under the conditions measured) or as unreliable because both have between-duplicate variance as a large fraction of the total.

### Identifying variable and stable sites on the Illumina EPIC array

To identify variably methylated and stably methylated probes on the Illumina EPIC array across individuals, we applied a downsampling approach. First, to ensure we detect robust variable and stable sites, in step 1, we downsampled the discovery data set by 10% (by removing randomly 10% of participants in the dataset). Second, to measure variability and stability, in step 2, we calculated standard deviation (SD) of methylation values for each probe across all individuals in our downsampled dataset. Those with the top 10% highest SD values were labelled as variable methylated probes (VMPs) and those with the 10% lowest SD values were labelled as the stable methylated probes (SMPs). We repeated the downsampling (step 1) and identification of VMPs and SMPs (step 2) ten times and kept only the VMPs and SMPs probes that appeared in all ten downsampled datasets. This analysis was then repeated in our validation dataset and only the VMPs and SMPs that were identified in both discovery and validation datasets were carried forward for further analysis. This provided us with a highly robust catalogue of CpG sites which had either variable (VMPs) or stable (SMPs) DNA methylation patterns (see Supplementary Files S1 and S2).

### Housekeeping and tissue specific gene annotations

We utilised expression data for 57 tissue types from the Epigenome Roadmap (73) and annotated housekeeping genes by assessing those genes where expression was in the top 40th percentile in all 57 samples, as previously described in (59). All other genes were then annotated as tissue specific genes.

### Genomic annotations and Gene ontology analyses

We annotated CpG sites using the manufacturer supplied annotation data (MethylationEPIC_v-1-0_B2 manifest file). Annotation was completed in the R package Minfi (74). Several categories were used as annotations in relation to CpG islands and divided into the following categories: CGIs, CGI shores (S and N), CGI shelfs (S and N) and open sea regions. Further, we also annotated the autosomal CpGs to several genomic features, including exons, introns, 5’ UTR, 3’UTR, enhancers, promoters and transposable elements (TEs) using data from UCSC table browser (https://genome.ucsc.edu/cgi-bin/hgTables). GO analyses were conducted using the gometh function in the missMethyl package (75), which tests gene ontology enrichment for significant CpGs while accounting for the differing number of probes per gene present on the EPIC array. For GO ontology analyses of enriched TFBS we used enrichGO and enrichKEGG from the clusterProfiler package in R (76), which performs false discovery rate (FDR) adjustment.

### Protein-protein network visualisation and hub gene identification

We searched all of the transcription factors enriched at VMPs and SMPs using the Search Tool for the Retrieval of Interacting Genes (STRING) (https://string-db.org) database to generate our TF networks. We extracted protein-protein interactions with a combined score greater than 0.4. Following this, we utilised the cytoscape plugin tool

Cytohubba (77) to characterise hub genes within the transcription factor networks. This was done by employing the local based method called maximum clique centrality (MCC).

### Enrichment of variably methylated probes and stably methylated probes in transcription factor binding motifs

The enrichment analysis of known motifs at VMPs and SMPs at promoters and enhancers was performed using the R package PWMEnrich (78) using the MotifDb collection of TF motifs (79). Specifically, the DNA sequences within a 50 bp range from the VMPs were extracted from the genome and compared to the SMPs as the background to reveal unique enriched motifs (adjusted p-value smaller than 0.05).

### Annotation of VMPs and SMPs to methylation quantitative trait loci

To investigate the proportion of VMPs and SMPs under genetic control, we annotated them to mQTLs using data from a previous study (24). We further characterised these into *cis, trans* and *cis* and *trans* mQTLs. We defined *cis* mQTLs as cases where the SNP and CpG site were located within 500 bp of each other and *trans* as cases where the SNP and the CpG site were greater 500 bp apart. Several CpG sites were annotated to more than one SNP, where one was a *cis* mQTL and another was a *trans* mQTL, thereby we classified these as *cis* and *trans* mQTLs. To annotate the mQTLs SNPs to biological processes we performed KEGG and GO enrichment analyses as previously described (see Genomic annotations and Gene ontology analyses).

### Chromatin loops and topologically associated domains

We examined whether the mQTL pairs occupied the same TAD or were connected by loops using Hi-C data available from the GEO under accession number (GSE124974) for white blood cells and neutrophils. Pre-processing of the data was performed using Juicer command line tools (74). Reads were aligned to the human (hg38) genome using BWA-mem (76) and then pre-processed using the Juicer pre-processing pipeline. We called chromatin loops using the HICCUPS tool from Juicer using a 10 Kb resolution. We also analysed whether our SNP and CpG pairs annotated by the mQTL analysis occupied the same topologically associated domain by using the hicFindTADs tool from HiCExplorer (80). These steps then allowed us to calculate the percentage of mQTL pairs which were connected via loops or occupying the same TAD compared to the EPIC array background. Fisher’s exact test was used to determine whether the differences were statistically significant. Lastly, Hi-C maps were generated using the hicPlotTADs function from HiCExplorer.

### Characterising putative epialleles

In order to characterise putative epialleles in human whole blood, we first identified those VMPs which had an average intermediate methylation across our sample as characterised by an average beta value between 0.40 and 0.60. Following this, we employed Hartigan’s dip test (81) using the diptest package in R to test for unimodality across these sites. We characterised epialleles as sites which had variable intermediate methylation and bimodal distribution (Hartigan’s dip test, p value smaller than 0.05); see Supplementary Files S3, S6 and S7.

### Integration with gene expression

We investigated the correlation between methylation variation and target gene expression variation at putative epialleles using gene expression microarray data obtained from the study of health in Pomerania (SHIP-Trend). This study is a longitudinal population based study based in Germany, aiming to assess common diseases and their relevant risk factors. Examinations took place from 2008-2012 and originally involved 4420 participants. Gene expression levels for a subset of these participants were measured using the Illumina HumanHT-12 v3 BeadChip array from whole blood cells (n=991). Details of the pre-processing methods have been previously reported in (82). We calculated the coefficient of variation for each gene as a measure of variance. The coefficient of variation was calculated by dividing the standard deviation of its expression value by its average expression value across the sample population. Following this, we then calculated Pearson’s correlation between methylation variation at each putative epiallele site and gene expression variation of their target gene.

## Availability of data and materials

The code to perform this analysis is available on GitHub https://github.com/livygrant97/interindividualVariation.

Supplementary File S1 - List of all SMPs in whole blood

Supplementary File S2 - List of all VMPs in whole blood

Supplementary File S3 - List of all putative epialleles in whole blood

Supplementary File S4 - List of all VMPs in brain

Supplementary File S5 - List of all VMPs in skeletal muscle

Supplementary File S6 - List of all putative epialleles in brain

Supplementary File S7 - List of all putative epialleles in skeletal muscle

## Author’s contributions

O.A.G., L.S. and N.R.Z. conceived and designed the experiments. O.A.G. performed the experiments and analysed the data. L.S., M.K. and N.R.Z. supervised the work. O.A.G., L.S., M.K. and N.R.Z. wrote the paper. The authors read and approved the final manuscript.

## Acknowledgements

The authors acknowledge the use of the High Performance Computing Facility at the University of Essex and would like to thank Stuart Newman for his support.

O.A.G. was funded by the University of Essex. M.K. was supported by the University of Essex and ESRC (grant RES-596-28-0001). L.S. was supported by Medical Research Council grant K013807. N.R.Z. was supported by the Queen Mary University of London. The analysis was facilitated by access to the Ceres high-performance computing cluster at the University of Essex.

