## Supplementary Material for "Systematic investigation of interindividual variation of DNA methylation in human whole blood"

Table S1: Enriched KEGG terms for VMPs and SMPs

**Enriched KEGG terms for SMPs**

| Description | N | DE | P.DE | FDR |
|---|---|---|---|---|
| Amyotrophic lateral sclerosis | 351 | 265 | 6,28122063667143e-15 | 2,21727088474501e-12 |
| Ribosome | 134 | 108 | 3,83876639434824e-14 | 6,77542268602464e-12 |
| Spliceosome | 132 | 112 | 7,91161858131672e-13 | 9,30933786401601e-11 |
| Parkinson disease | 252 | 193 | 1,47771751366563e-12 | 1,30408570580992e-10 |
| Cell cycle | 127 | 110 | 2,95604276231233e-12 | 2,0869661901925e-10 |
| Pathways of neurodegeneration - multiple diseases | 462 | 333 | 8,10586829637703e-12 | 4,76895251436849e-10 |
| Alzheimer disease | 370 | 271 | 1,50070757741972e-11 | 7,56785392613087e-10 |
| Huntington disease | 292 | 217 | 3,51626417177965e-11 | 1,55155156579777e-09 |
| Shigellosis | 244 | 188 | 5,79018166509307e-11 | 2,27103791975317e-09 |
| Protein processing in endoplasmic reticulum | 169 | 137 | 7,54673414710612e-11 | 2,66399715392846e-09 |
| mTOR signaling pathway | 155 | 129 | 8,78760976760718e-11 | 2,82002386178667e-09 |
| Oxidative phosphorylation | 121 | 97 | 2,2300125731345e-10 | 6,55995365263732e-09 |
| Thermogenesis | 219 | 165 | 3,40217144992078e-10 | 9,23820401401567e-09 |
| Nucleocytoplasmic transport | 107 | 91 | 4,1923778216466e-10 | 1,05707812217232e-08 |
| Metabolic pathways | 1520 | 960 | 5,4918214935511e-10 | 1,29240865814902e-08 |
| Prion disease | 259 | 188 | 1,95789911948829e-09 | 4,31961493237104e-08 |
| Ubiquitin mediated proteolysis | 142 | 117 | 2,72565075958291e-09 | 5,06397220069877e-08 |
| Endocytosis | 250 | 193 | 2,67319877609672e-09 | 5,06397220069877e-08 |
| Human papillomavirus infection | 330 | 243 | 2,58302454061205e-09 | 5,06397220069877e-08 |
| Salmonella infection | 248 | 186 | 9,65910682471857e-09 | 1,70483235456283e-07 |
| Lysosome | 132 | 107 | 1,45490779197195e-08 | 2,44563071698142e-07 |
| p53 signaling pathway | 73 | 65 | 2,07368240990917e-08 | 3,32731768499062e-07 |
| RNA degradation | 79 | 67 | 3,64316881515636e-08 | 5,59147213804433e-07 |
| Non-alcoholic fatty liver disease | 151 | 115 | 6,07384643024182e-08 | 8,93361579114735e-07 |
| Aminoacyl-tRNA biosynthesis | 43 | 41 | 6,62322928392709e-08 | 9,35199974890504e-07 |
| Cellular senescence | 155 | 123 | 7,03241228125542e-08 | 9,5478520587814e-07 |
| Hepatocellular carcinoma | 168 | 132 | 7,56743751176594e-08 | 9,89372385797547e-07 |
| Chronic myeloid leukemia | 76 | 67 | 1,45689697077046e-07 | 1,83673082386419e-06 |
| mRNA surveillance pathway | 95 | 77 | 1,65484563259968e-07 | 2,01434658037133e-06 |
| Proteasome | 46 | 42 | 1,7525083919525e-07 | 2,0621182078641e-06 |
| Ribosome biogenesis in eukaryotes | 76 | 62 | 2,25833106396968e-07 | 2,57158343735902e-06 |
| Proteoglycans in cancer | 204 | 155 | 4,38633309553444e-07 | 4,83867369601143e-06 |
| Diabetic cardiomyopathy | 189 | 137 | 7,35093955530803e-07 | 7,86327776673859e-06 |
| Viral carcinogenesis | 193 | 141 | 1,19313842083981e-06 | 1,23875841928369e-05 |
| Autophagy - animal | 140 | 109 | 3,23573425894806e-06 | 3,26346912402475e-05 |
| Neurotrophin signaling pathway | 119 | 95 | 3,34862279106797e-06 | 3,28351068124165e-05 |
| Pathways in cancer | 529 | 355 | 9,22607496892237e-06 | 8,80217422710701e-05 |
| Insulin signaling pathway | 137 | 105 | 1,22318057380546e-05 | 0,000113627037514034 |

| | | | | |
|---|---|---|---|---|
| Hepatitis B | 162 | 117 | 2,0755261774501e-05 | 0,000187861728369201 |
| Protein export | 23 | 22 | 2,26024468994379e-05 | 0,000199466593887539 |
| Bacterial invasion of epithelial cells | 77 | 64 | 2,33406429394967e-05 | 0,000200957242869325 |
| Pancreatic cancer | 76 | 63 | 2,82469897306439e-05 | 0,000237409223212316 |
| Basal cell carcinoma | 63 | 53 | 2,92865109557689e-05 | 0,000240421822497358 |
| AMPK signaling pathway | 120 | 93 | 3,36940611651206e-05 | 0,000268453281014652 |
| Pathogenic Escherichia coli infection | 196 | 140 | 3,42220896477602e-05 | 0,000268453281014652 |
| Colorectal cancer | 86 | 69 | 4,37258388979409e-05 | 0,000335548285455938 |
| Yersinia infection | 135 | 102 | 4,99864012234574e-05 | 0,000375429779401712 |
| Chemical carcinogenesis - reactive oxygen species | 209 | 143 | 5,33989998021755e-05 | 0,000392705144378499 |
| Longevity regulating pathway | 89 | 71 | 5,5421314174805e-05 | 0,000399259671504208 |
| Vibrio cholerae infection | 50 | 42 | 7,35047831314703e-05 | 0,000508768400890373 |
| Human cytomegalovirus infection | 225 | 154 | 7,24606929402117e-05 | 0,000508768400890373 |
| EGFR tyrosine kinase inhibitor resistance | 78 | 64 | 8,74667469087359e-05 | 0,000593764647284303 |
| Epstein-Barr virus infection | 199 | 139 | 0,000102372493468261 | 0,000681839437628229 |
| Base excision repair | 33 | 29 | 0,000107771033517345 | 0,000704503237622645 |
| Prostate cancer | 97 | 76 | 0,00011431415727259 | 0,000733689045767715 |
| Viral life cycle - HIV-1 | 63 | 50 | 0,000123251924138016 | 0,000776927307512851 |
| Human immunodeficiency virus 1 infection | 211 | 143 | 0,000129305197858292 | 0,000800784821824158 |
| Non-small cell lung cancer | 72 | 59 | 0,000135720973630878 | 0,000826025925718964 |
| Apoptosis | 135 | 99 | 0,000143715595053966 | 0,000859857712780506 |
| Necroptosis | 157 | 104 | 0,000167335341466477 | 0,00098448959229444 |
| Hippo signaling pathway | 157 | 117 | 0,000179647339085533 | 0,00103959853601956 |
| Notch signaling pathway | 58 | 48 | 0,00019787611707035 | 0,00112661724719086 |
| Coronavirus disease - COVID-19 | 231 | 142 | 0,000213466251292555 | 0,00119351054566879 |
| Glioma | 75 | 60 | 0,000216387181084426 | 0,00119351054566879 |
| Spinocerebellar ataxia | 142 | 105 | 0,000253491517351228 | 0,00137665393269206 |
| Endocrine resistance | 96 | 75 | 0,000266263901250229 | 0,00142410844153531 |
| N-Glycan biosynthesis | 50 | 41 | 0,000303622142344774 | 0,00157615612128978 |
| Insulin resistance | 108 | 81 | 0,000301393919488287 | 0,00157615612128978 |
| Breast cancer | 147 | 108 | 0,000316740602661474 | 0,00162042656144203 |
| Human T-cell leukemia virus 1 infection | 219 | 154 | 0,000367491521422416 | 0,00185320724374447 |
| Renal cell carcinoma | 68 | 55 | 0,000386847161957786 | 0,00192333870663519 |
| Cushing syndrome | 155 | 113 | 0,000466201088024551 | 0,00228568033434259 |
| FoxO signaling pathway | 131 | 95 | 0,000508173248435117 | 0,0024573309136657 |
| Gastric cancer | 149 | 108 | 0,000548927288692065 | 0,00261853152578782 |
| Various types of N-glycan biosynthesis | 39 | 33 | 0,000669719262003778 | 0,00315214532649778 |
| DNA replication | 36 | 30 | 0,000763439097795538 | 0,00353583605703785 |
| Hepatitis C | 157 | 106 | 0,000771273020940267 | 0,00353583605703785 |
| Hedgehog signaling pathway | 56 | 46 | 0,000821624312240712 | 0,00371837669514066 |
| Kaposi sarcoma-associated herpesvirus infection | 194 | 129 | 0,000886477288587348 | 0,0039610947198903 |

| | | | | |
|---|---|---|---|---|
| Tight junction | 169 | 118 | 0,00102978025007159 | 0,00448780775648484 |
| Endometrial cancer | 58 | 47 | 0,00102958143960272 | 0,00448780775648484 |
| Adherens junction | 71 | 56 | 0,00110537116140705 | 0,00470115686718903 |
| Small cell lung cancer | 92 | 70 | 0,00110152679335838 | 0,00470115686718903 |
| PD-L1 expression and PD-1 checkpoint pathway in cancer | 89 | 66 | 0,00150752709905882 | 0,00633520316628289 |
| Mitophagy - animal | 71 | 54 | 0,00154180161208694 | 0,00640301140078458 |
| Oocyte meiosis | 126 | 88 | 0,00172246532470681 | 0,00707011929792446 |
| Sphingolipid signaling pathway | 118 | 86 | 0,00200549545610396 | 0,00813724018396206 |
| Focal adhesion | 200 | 142 | 0,00204912873718499 | 0,00821980050257159 |
| Terpenoid backbone biosynthesis | 23 | 20 | 0,00249490928256442 | 0,00989553906455327 |
| Phosphatidylinositol signaling system | 97 | 72 | 0,00314494744729239 | 0,0123351827654913 |
| Epithelial cell signaling in Helicobacter pylori infection | 70 | 52 | 0,00328711781293906 | 0,0127511273403021 |
| Regulation of actin cytoskeleton | 228 | 155 | 0,00370379827737956 | 0,0142113129555977 |
| RNA polymerase | 34 | 25 | 0,00388269689833619 | 0,0146706444756503 |
| MAPK signaling pathway | 294 | 198 | 0,00390663054025816 | 0,0146706444756503 |
| TNF signaling pathway | 112 | 78 | 0,00397285117408299 | 0,0147622785731715 |
| Signaling pathways regulating pluripotency of stem cells | 142 | 100 | 0,0042084709121646 | 0,0154748982499386 |
| Thyroid hormone signaling pathway | 121 | 88 | 0,00481004263864616 | 0,0175045881591969 |
| Thyroid cancer | 37 | 30 | 0,00509737708914352 | 0,0183609603313027 |
| Choline metabolism in cancer | 98 | 72 | 0,00535996099187389 | 0,0191117801023382 |
| Other glycan degradation | 18 | 16 | 0,00603438399320163 | 0,0213013754960018 |
| Fc gamma R-mediated phagocytosis | 96 | 69 | 0,00621317655651187 | 0,0217153596480068 |
| Ferroptosis | 41 | 32 | 0,00638083588192775 | 0,0220826967286323 |
| Fanconi anemia pathway | 53 | 39 | 0,0068124578658109 | 0,0233475497731189 |
| Nucleotide excision repair | 46 | 34 | 0,00827867615657796 | 0,028099737339154 |
| Growth hormone synthesis, secretion and action | 120 | 85 | 0,00868533398513894 | 0,0291992656833719 |
| Selenocompound metabolism | 17 | 15 | 0,00884730238042067 | 0,0294631862291368 |
| Parathyroid hormone synthesis, secretion and action | 106 | 76 | 0,00933748154921164 | 0,0308049624941281 |
| Lysine degradation | 63 | 47 | 0,00955265201093819 | 0,0312230199987146 |
| Basal transcription factors | 44 | 32 | 0,0104113051102941 | 0,0337173459076497 |
| Non-homologous end-joining | 13 | 12 | 0,0106399110727049 | 0,034144441896953 |
| Valine, leucine and isoleucine degradation | 48 | 36 | 0,0112333783321689 | 0,0357241671284289 |
| Prolactin signaling pathway | 70 | 51 | 0,0119061030084194 | 0,0375254853747505 |
| One carbon pool by folate | 20 | 17 | 0,0121476550317687 | 0,0379479843027819 |
| Melanogenesis | 101 | 71 | 0,0125586623645185 | 0,0388877878480267 |
| Wnt signaling pathway | 169 | 116 | 0,0129618352621462 | 0,0397871986742402 |
| Glycosylphosphatidylinositol (GPI)-anchor biosynthesis | 26 | 20 | 0,0139020096461422 | 0,0409728164774185 |
| Platinum drug resistance | 73 | 51 | 0,0137100191288479 | 0,0409728164774185 |
| HIF-1 signaling pathway | 109 | 75 | 0,0136467688844695 | 0,0409728164774185 |
| VEGF signaling pathway | 59 | 44 | 0,0139284361962896 | 0,0409728164774185 |
| Glucagon signaling pathway | 105 | 71 | 0,0139079206134487 | 0,0409728164774185 |

| | | | | |
|---|---|---|---|---|
| Central carbon metabolism in cancer | 70 | 51 | 0,0141732165106753 | 0,0413483093245321 |
| Measles | 139 | 88 | 0,0149435620874029 | 0,0432383394824035 |

**Enriched KEGG terms for VMPs**

| Description | N | DE | P.DE | FDR |
|---|---|---|---|---|
| Olfactory transduction | 407 | 162 | 1.76027586223357e-09 | 6.21377379368449e-07 |
| Neuroactive ligand-receptor interaction | 363 | 195 | 3.53430651674902e-06 | 0.000623805100206203 |

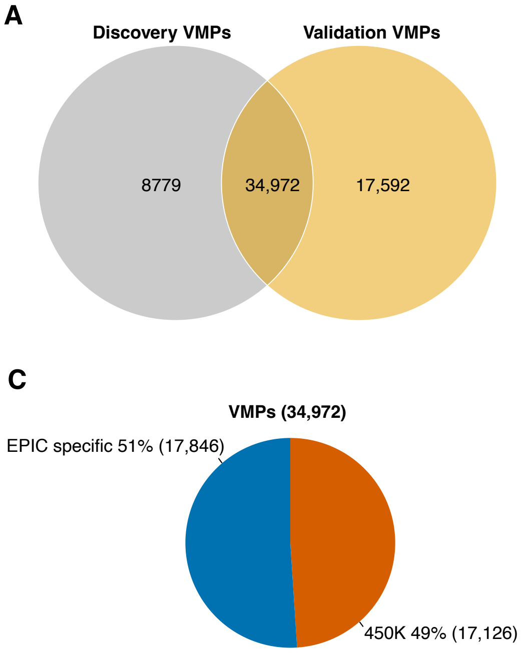


Figure S1: *Identification of VMPs and SMPs.* Venn diagram showing the number of VMPs (A) and SMPs (B) identified in discovery and validation data sets. The percentage of VMPs (C) and SMPs (D) that are EPIC specific probes.

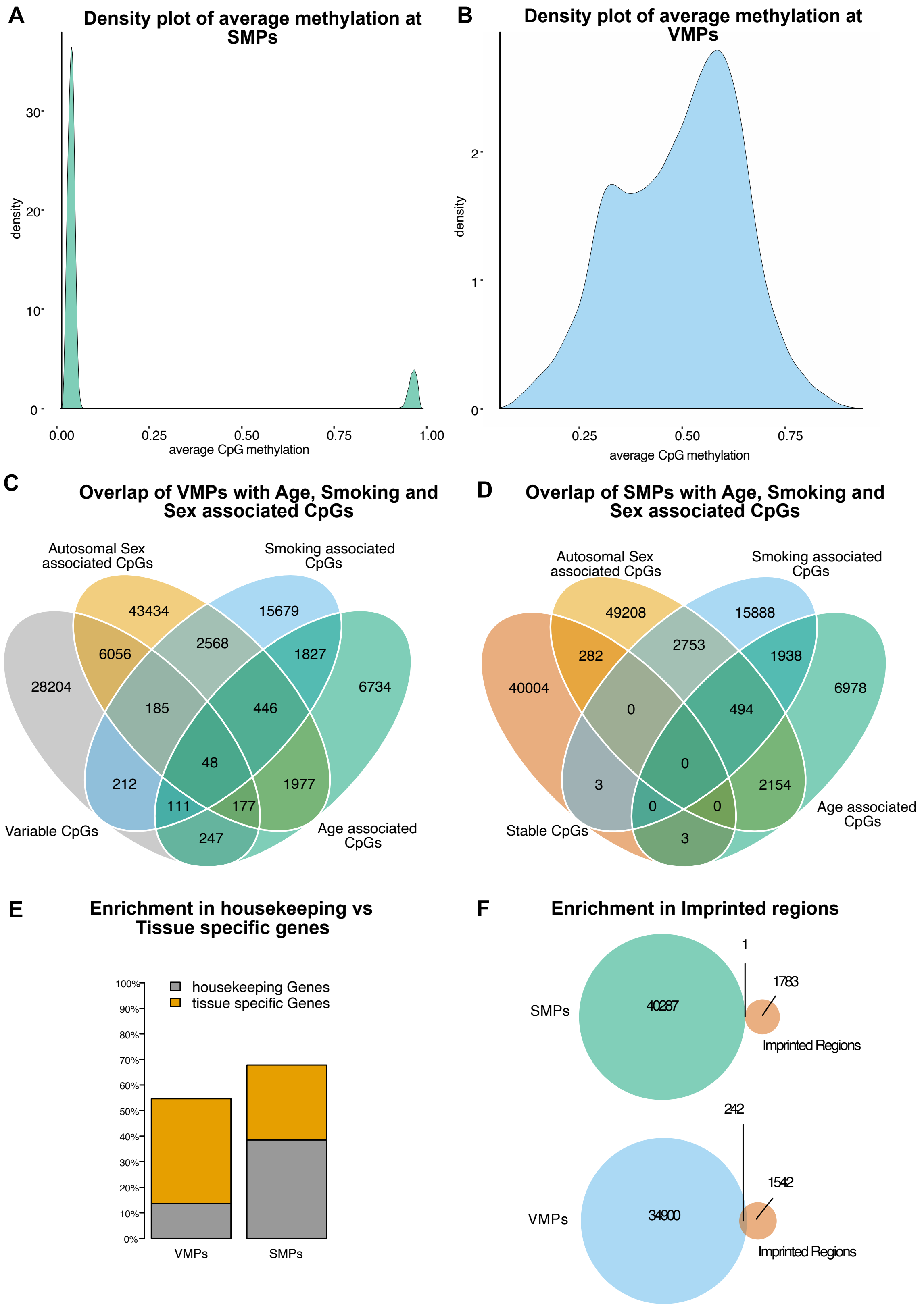


Figure S2: *Characterisation of VMPs and SMPs.* Density plots showing distribution of average CpG methylation at (A) SMPs and (B) VMPs. Venn diagrams showing overlap of (C) VMPs and (D) SMPs with known autosomal sex, smoking and age associated CpGs. (E) Barplot representing the enrichment of VMPs and SMPs in housekeeping and tissue specific genes. (F) Venn diagrams representing enrichment of SMPs and VMPs in imprinted regions.

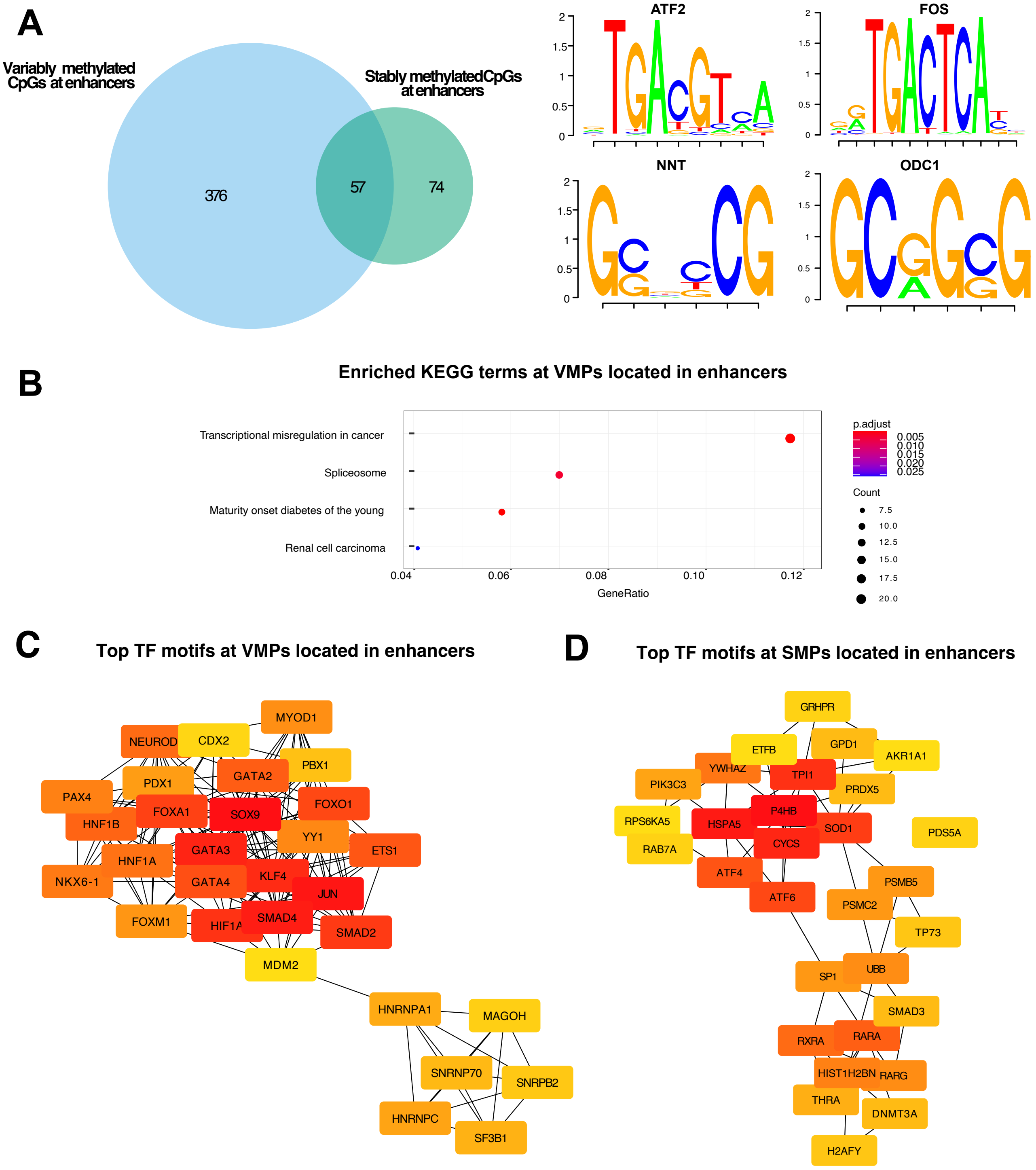


Figure S3: *Transcription factor motif enrichment for VMPs and SMPs at promoters.* (A) Overlap of enriched TF motifs for VMPs (blue) and SMPs (green). The top two motifs enriched in VMPs were ATF2 and FOS and in SMPs were NNT and ODC1. (B) KEGG analyses for the significantly enriched TF motifs at VMPs and SMPs at enhancers. (C-D) Sub networks of the top 30 enriched TF motifs at (C) VMPs at enhancers and (D) SMPs at enhancers. Node colour represents the degree of connectivity. The scale from red to yellow represents the top 30 enriched TF motif rank from 1-30, with red indicating highest degree and yellow indicating lowest degree.

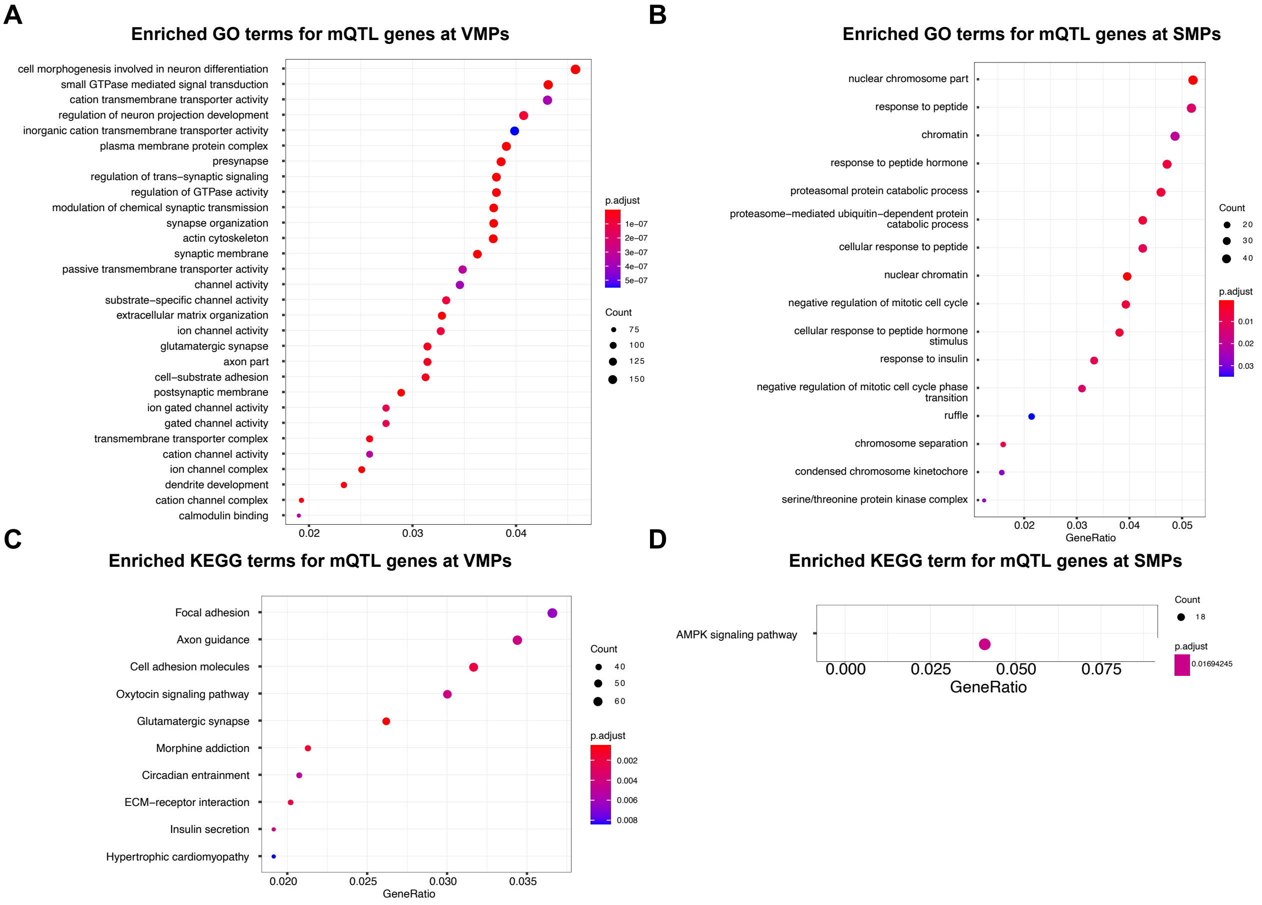


Figure S4: *Enriched GO and KEGG terms for mQTLs.* (A) GO and (C) KEGG analyses for the mQTL genes at VMPs. (B) GO and (D) KEGG analyses for the mQTL genes at SMPs.

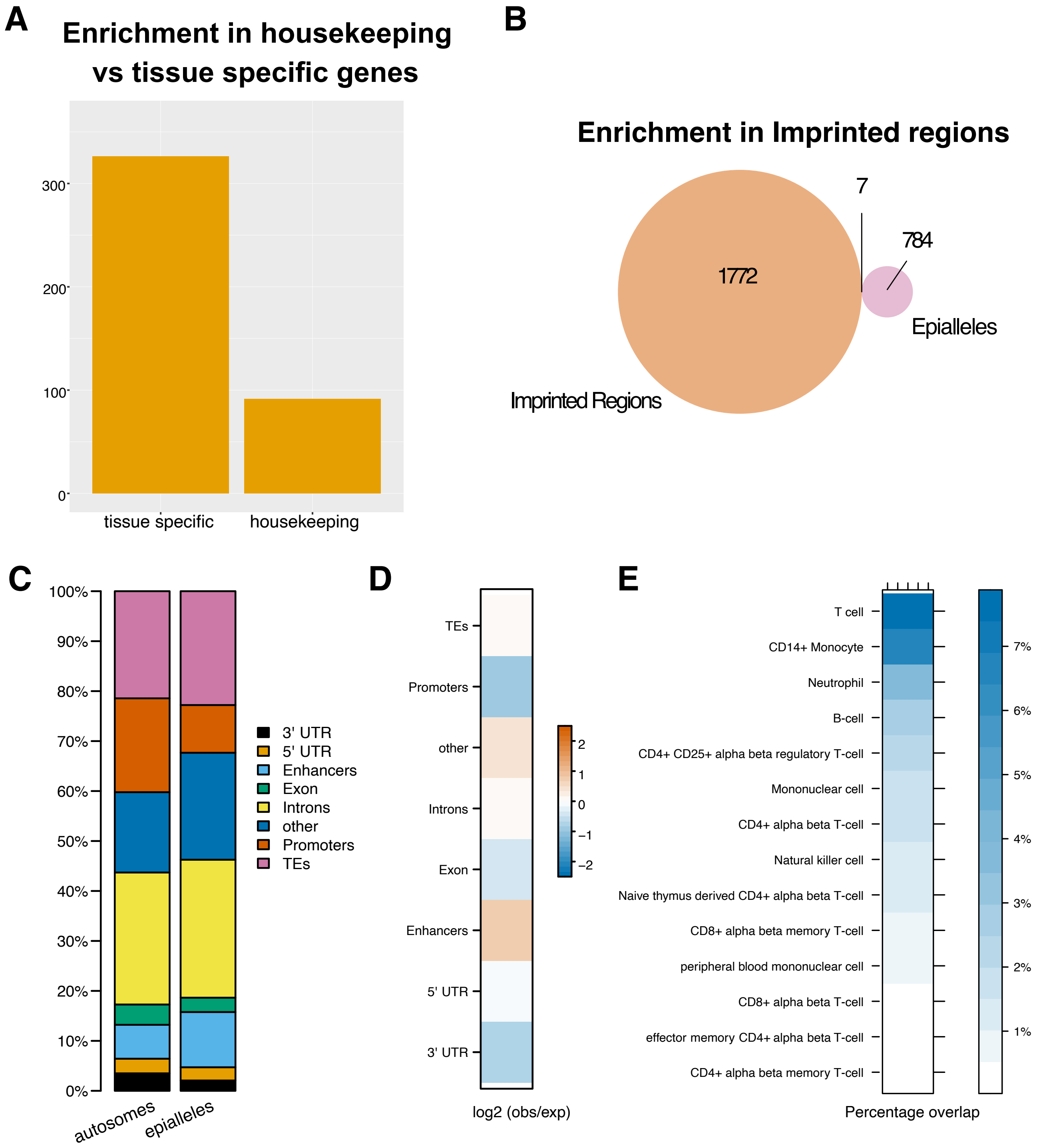


Figure S5: *Functional annotations of putative epialleles in human whole blood.* (A) The overlap between putative epialleles and housekeeping or tissue specific genes. (B) The overlap between putative epialleles and imprinted loci. (C) The overlap of all putative epialleles (n=784) with genomic features compared to the background. (D) The log2 (obs/exp) based on the background of the different annotations. (E) The percentage of epialleles that overlap with H3K27me3 peaks in several blood cell lines.

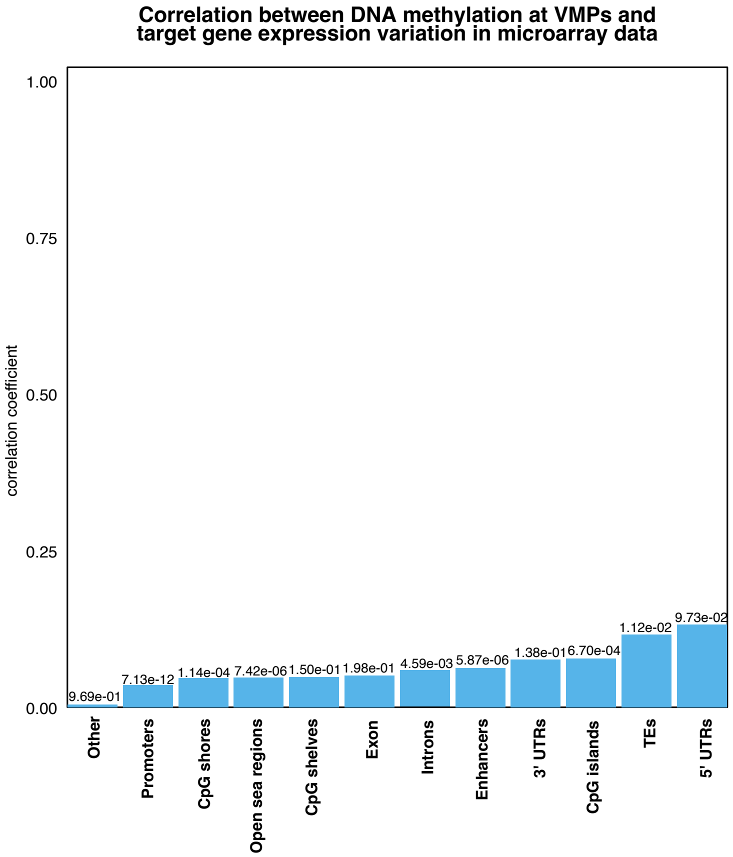


Figure S6: *Link between methylation and expression at VMPs.* The correlation between variation in gene expression and variation in methylation of VMPs. The correlation test significance is reported above the bars.

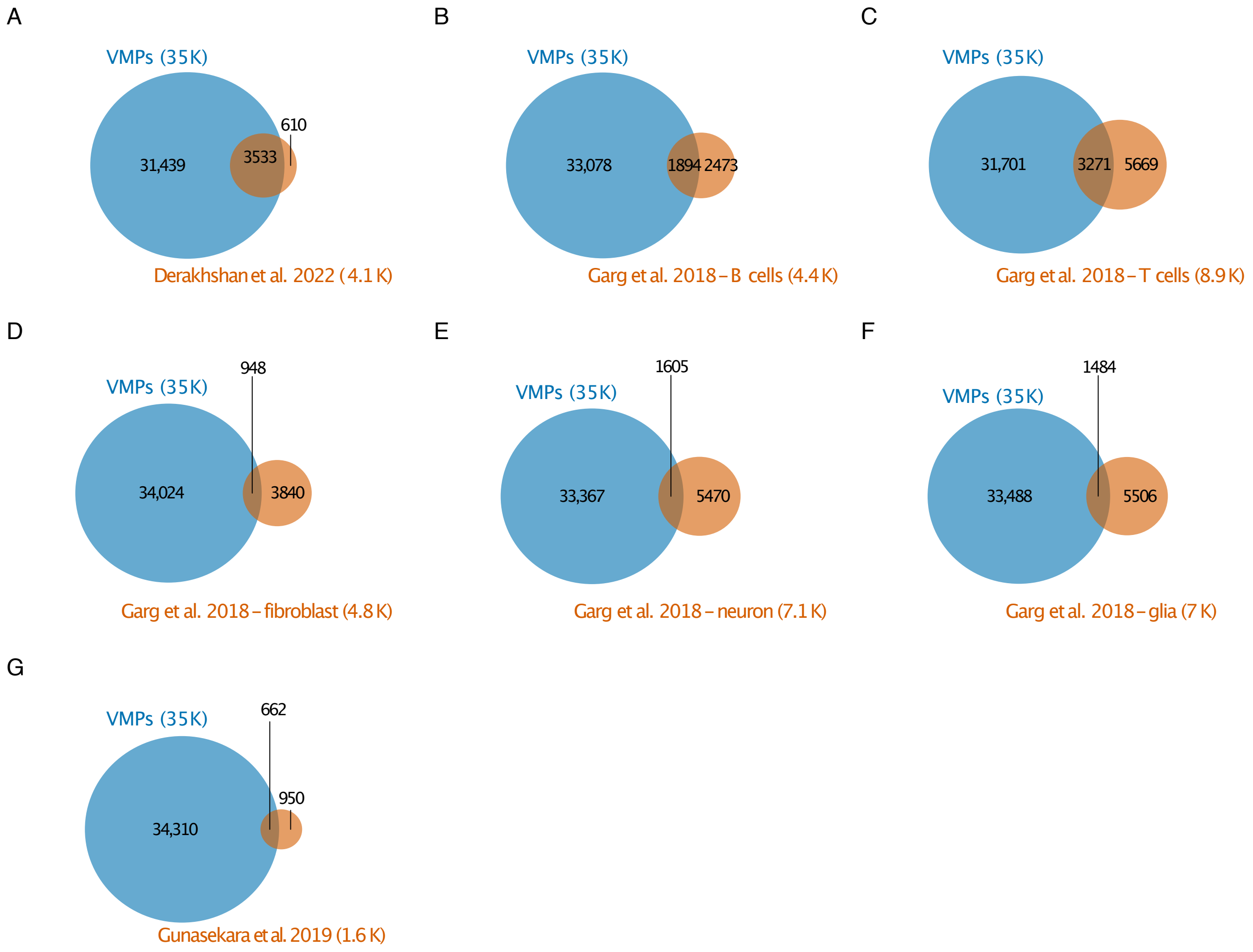


Figure S7: *Overlap of VMPs with CpGs identified to display variable interindividual methylation in other studies.* Venn diagram showing the number of VMPs that overlap with variable probes identified in previous studies: (A) multiple tissues in (Derakhshan et al. 2022), (B) B cells in (Garg et al. 2018), (C) T cells in (Garg et al. 2018), (D) fibroblast cells in (Garg et al. 2018), (E) neurons in (Garg et al. 2018), (F) glia in (Garg et al. 2018) and (G) multiple tissues in (Gunasekara et al. 2019). In the latter study, the authors identified regions displaying variable DNA methylation using bisulfite sequencing and to keep make the studies comparable, we selected only the CpGs on the EPIC array that were located in those regions.
